# GATA2 Induces a Stem Cell-Like Transcriptional Program in Macrophages that Promotes Atherogenesis

**DOI:** 10.1101/2025.07.18.665581

**Authors:** Amena Aktar, Angela M Vrieze, Kiera Telesnicki, Paisley Cox-Duvall, Matthew Arbolino, Rodney P DeKoter, A. Dave Nagpal, Bryan Heit

## Abstract

Atherosclerosis is a chronic inflammatory disease characterized by the accumulation of lipid-laden necrotic macrophages within blood vessels walls. GATA2 is a normally hematopoietic transcription factor which in the bone marrow helps maintain the proliferative, non-differentiated phenotype of hematopoietic progenitors. Unexpectedly, GATA2 is upregulated in macrophages within atherosclerotic plaque, where it plays an unknown role in disease progression. Although GATA2 can be expressed from two promoters, we determined that the atherogenic stimuli oxidized low-density lipoprotein and TNFα induce GATA2 expression via the internal (IG) GATA2 promoter, with GATA2 transcription initiated by the transcription factors NF-κB, STAT1, and the aryl hydrocarbon receptor. GATA2 had a divergent effect on promoter activity, with GATA2 upregulating genes associated with stem cell maintenance, proliferation, reactive oxygen species production, and migration, while downregulating genes central to macrophage function including those for cholesterol efflux, pathogen phagocytosis, and for the efferocytosis of apoptotic cells. Consequentially, GATA2-expressing macrophages had a pro-atherogenic phenotype typified by an invasive phenotype, poor cholesterol efflux, and impaired phagocytosis and efferocytosis. These results indicate that GATA2 upregulation induces an immature, stem cell-like phenotype in atheroma macrophages, thereby promoting plaque cellularity while compromising atheroprotective mechanisms such as cholesterol clearance and apoptotic cell removal.

## Introduction

Macrophages are the primary cell type that mediate atherosclerosis, where the accumulation of lipids, release of inflammatory cytokines, and eventual necrosis of these cells forms the lipid- and cell-debris-rich environment that typifies an atherosclerotic plaque^1,2^. The formation of these lesions is initiated by the deposition and oxidation of low-density lipoprotein (LDL) within the vascular intima^3^. Macrophages endocytose these deposits, and under non-disease conditions return the cholesterol to the circulation by exporting it to high density lipoprotein (HDL)^4^. Excess LDL deposition can overwhelm this system, leading to the induction of macrophage cell stress, driving reactive oxygen species production and the oxidation of LDL to form oxidized LDL (oxLDL). Unlike LDL, oxLDL is an inflammatory stimulus, inducing macrophage activation via signaling through the receptors CD36 and toll-like receptors 2 and 4^5^. This induces an inflammatory response in the intimal macrophages, resulting in the recruitment of monocytes which undergo local proliferation and differentiation into macrophages, further seeding the developing atheroma with inflammatory macrophages^6^. Atheroma macrophages undergo turnover in as little as four weeks, with the maintenance and growth of the macrophage population mediated primarily by monocyte recruitment in early atheromas but shifting almost entirely to local proliferation at later stages of disease^6,7^. Local signals within the atheroma further polarizes these macrophages into atherogenic and anti-atherogenic polarization states – for example, exposure to heme mediates formation of the Mheme phenotype, which exerts anti-inflammatory and anti-atherogenic effects on the plaque through the removal and metabolism of free heme^8^. These distinct macrophage subsets arise from polarization state-specific transcriptional programs, mediated by the selective upregulation of a small number of transcription factors.

We identified GATA2 as a transcription factor that is upregulated in a large percentage of macrophages in early-stage human atheromas. Here, GATA2 expression induces a proatherogenic phenotype typified by an inability to clear dying cells and cell debris from the forming lesion^9^. More recently we determined that this GATA2+ population persists into later stages of disease, where GATA2 expression drives the proliferation of atheroma macrophages^10^. Consistent with our observations, other studies have identified single nucleotide polymorphisms in GATA2 that predispose patients to atherosclerosis, but the mechanisms by which GATA2 drives disease remains unknown^11,12^. The expression of GATA2 in atheroma macrophages is unusual, as GATA2 is required for the proliferation of hematopoietic stem cells in the bone marrow, and for the commitment of hematopoietic stem cells to the myeloid lineage^13–16^. However, GATA2 expression must cease for myeloid progenitors to complete their maturation into monocytes and other myeloid cells. Thus, GATA2 expression is not typically observed in myeloid cells outside of the bone marrow, with the exception of some forms of myeloid leukemia where GATA2 is re-expressed and functions as an oncogene^14,17,18^. Consequentially, it is unknown what signals cause GATA2 to be upregulated in atheroma macrophages, which transcription factors mediate this upregulation, nor is it known what the direct impact of GATA2 expression is on macrophage gene expression.

As a member of the GATA family of transcription factors, GATA2 binds the DNA motif G-A-T-A via a tandem zinc finger domain^19^. GATA2 is a pioneering transcription factor that can bind to and open heterochromatin, thereby allowing other transcription factors access to the DNA^20^. While not previously investigated in macrophages, GATA2 is known to bind to and activate enhancer regions in mast cells, while in endothelial cells GATA2 was found to bind near AP-1 binding sites where GATA2 was required for maximal AP-1 activity^21,22^. In the bone marrow GATA2 serves two functions. During mitosis GATA2 “bookmarks” genes, allowing these genes to be readily expressed once DNA decondenses after cells enter interphase^23^. Secondly, GATA2 governs cell-fate decisions during hematopoiesis, often through competitive promoter binding with other GATA family transcription factors that exert opposing regulatory effects on target genes^24,25^. Furthermore, GATA2 activity can be posttranscriptionally regulated, with phosphorylation, SUMOylation, acetylation, and ubiquitination altering GATA2 nuclear localization, DNA binding affinity, and stability^26^. Thus, the impact of GATA2 on a cells transcriptional profile can be difficult to predict, as competitive and cooperative interactions with other transcription factors, enhancer versus core promoter binding, GATA2 expression level, and post-translational modifications can alter both GATA2 binding patterns and activity.

Given that GATA2 upregulation has only been reported in atheroma macrophages, and not in other human macrophage populations, we used a combination of cell-based assays and ChIP-seq to test the hypothesis that atheroma-specific inflammatory factors upregulate GATA2 in atheroma macrophages, resulting in suppressed expression of anti-atherogenic genes in these cells. These experiments revealed the transcription factors responsible for upregulating GATA2 in response to oxLDL and the pro-inflammatory cytokine TNFα. They also uncovered a GATA2-driven stem cell-like transcriptional program that disrupts cholesterol metabolism and reshapes the inflammatory landscape of these cells.

## Results

### CHARACTERIZATION OF THE MACROPHAGE GATA2 PROMOTER

GATA2 can be expressed from one of two promoters – the IS promoter located 5’ to the first exon of the GATA2 gene, and the IG promoter located between the first and second exons of GATA2 (**Figure 1A**)^27^. As the promoter controlling GATA2 expression in macrophages is unknown, we stimulated macrophages with oxLDL to induce GATA2 expression and then used 5’ RACE to identify the active promoter. This produced a PCR product of ∼550 bp, the size expected of the IG promoter, and larger than the ∼350 bp band expected of the IS promoter (**Figure 1B**). Sequencing of the PCR product and alignment against the GRCh38/hg38 human genome confirmed that the IG promoter was driving GATA2 expression in macrophages and identified the transcriptional start site (**Figure 1C**). The ENCODE ChIP database was used to identify putative transcription factor binding regions in the IG promoter, identifying sites for NF-κB, AP-1, the aryl hydrocarbon receptor (AHR), STAT1, and RORα (**Figure 1C**)^28^. Also identified in this region is a sequence competitively bound by GATA1 and GATA2, which is part of the GATA-switch mechanism used to inactivate GATA2 in the presence of active GATA1^24^. A dual-luciferase vector system containing firefly luciferase under control of the IG promoter was then assembled and transduced into THP-1-derived macrophages, where a low-level of basal GATA2 IG promoter activity was observed (**Figure 1D**). GATA2 IG promoter activity increased significantly when these macrophages were stimulated with oxLDL or TNFα (**Figure 1E**), indicating that GATA2 expression is induced in macrophages by plaque-resident lipoproteins and cytokines.

**Figure 1:**
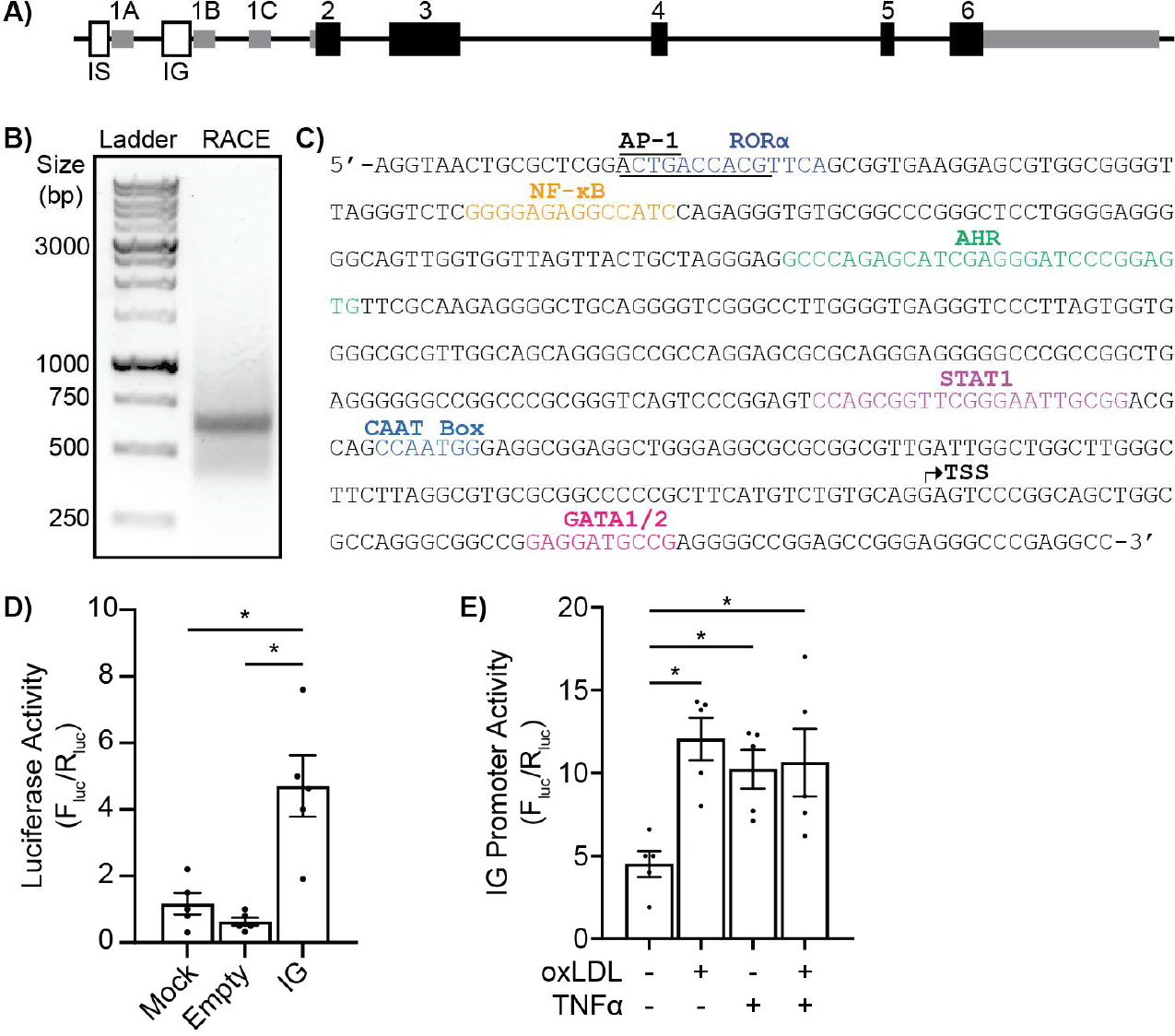
Identification and Characterization of the Macrophage GATA2 Promoter. **A)** Diagram of the GATA2 promoter. Exons are numbered and are indicated by solid boxes; non-coding regions are in grey and coding regions are in black. The location of the IS and IG promoter are indicated by white boxes. **B)** DNA gel showing the 5’ RACE product of the GATA2 mRNA amplified from THP1-derived macrophages stimulated with 100 μg/mL oxLDL. **C)** GATA2 promoter region identified by 5’ RACE, showing the transcriptional start site (TSS), CAAT box, and putative binding sites for AP-1, RORα, NF-κB, AHR, STAT1, and a GATA1/2 switch motif. **D)** Basal activity of the GATA2 IS promoter in THP1-derived macrophages measured using a dual-luciferase assay in which promotor activity is measured as the ratio of firefly luciferase activity (F_luc_), under control of the GATA2 IG promoter, normalized to Renilla luciferase activity (R_luc_), under control of a constitutively active promoter. Mock = mock-transfected cells, Empty = luciferase activity from a promoter-less F_luc_ vector, IG = F_luc_ under control of the GATA2 IG promoter. **E)** Dual-luciferase quantification of GATA2 promoter activity in THP1 macrophages stimulated for 48 hours with 100 μg/mL oxLDL or 10 ng/mL TNFα. n = 5 biological replicates, * p < 0.05 between the indicated groups, ANOVA with Tukey correction.

Given the presence of NF-κB, AP-1, AHR, STAT1, and RORα binding motifs in the GATA2 IG promoter, we next investigated the role of these transcription factors in mediating IG promoter activity in response to oxLDL and TNFα. Inhibition of NF-κB with BMS-345541 inhibited GATA2 IG promoter activity in resting macrophages, and suppressed both oxLDL- and TNFα-induced increases in GATA2 IG promoter activity (**Figure 2A**). Both the STAT1 inhibitor nifuroxazide (**Figure 2B**) and the AHR inhibitor CH223191 (**Figure 2C**) inhibited oxLDL- and TNFα-induced increases in GATA2 IG promoter activity, but did not affect basal GATA2 promoter activity. These data are consistent with GATA2 expression being induced by atherogenic stimuli^5,29^. Interestingly, activation of RORα with all-trans retinoic acid (AtRA) increased basal GATA2 IG promoter activity while suppressing TNFα-induced promoter activity (**Figure 2D**), whereas inhibition of AP-1 with SR11302 enhanced both basal GATA2 IG promoter activity and promoter activity in response to simultaneous treatment with oxLDL and TNFα (**Figure 2E**), indicating that RORα and AP-1 negatively regulate GATA2 IG in response to inflammatory stimuli. Because pharmacological inhibitors can have off-target effects, we generated mutants of the IG promoter luciferase reporter lacking the NF-κB or STAT1 binding sites (**Figure 1C**). Deletion of the NF-κB site reduced IG promoter activity in response to both oxLDL and TNFα, whereas deletion of the STAT1 binding site only reduced IG promoter activity in response to TNFα (**Figure 2F**). Both oxLDL and TNFα are known to be potent inducers of NF-κB activity, but neither have been reported to activate STAT1. Given these assays were performed over 48 hours, the STAT1 activity may have been induced by autocrine or paracrine signalling downstream of a cytokine produced in response to TNFα stimulation. In macrophages TNFα induces, and oxLDL suppresses, IL-1β secretion—a potent activator of STAT1—potentially explaining the differential impact of NFκB versus STAT1 binding site deletion on the activity of the GATA2 IG promoter^30^.

**Figure 2:**
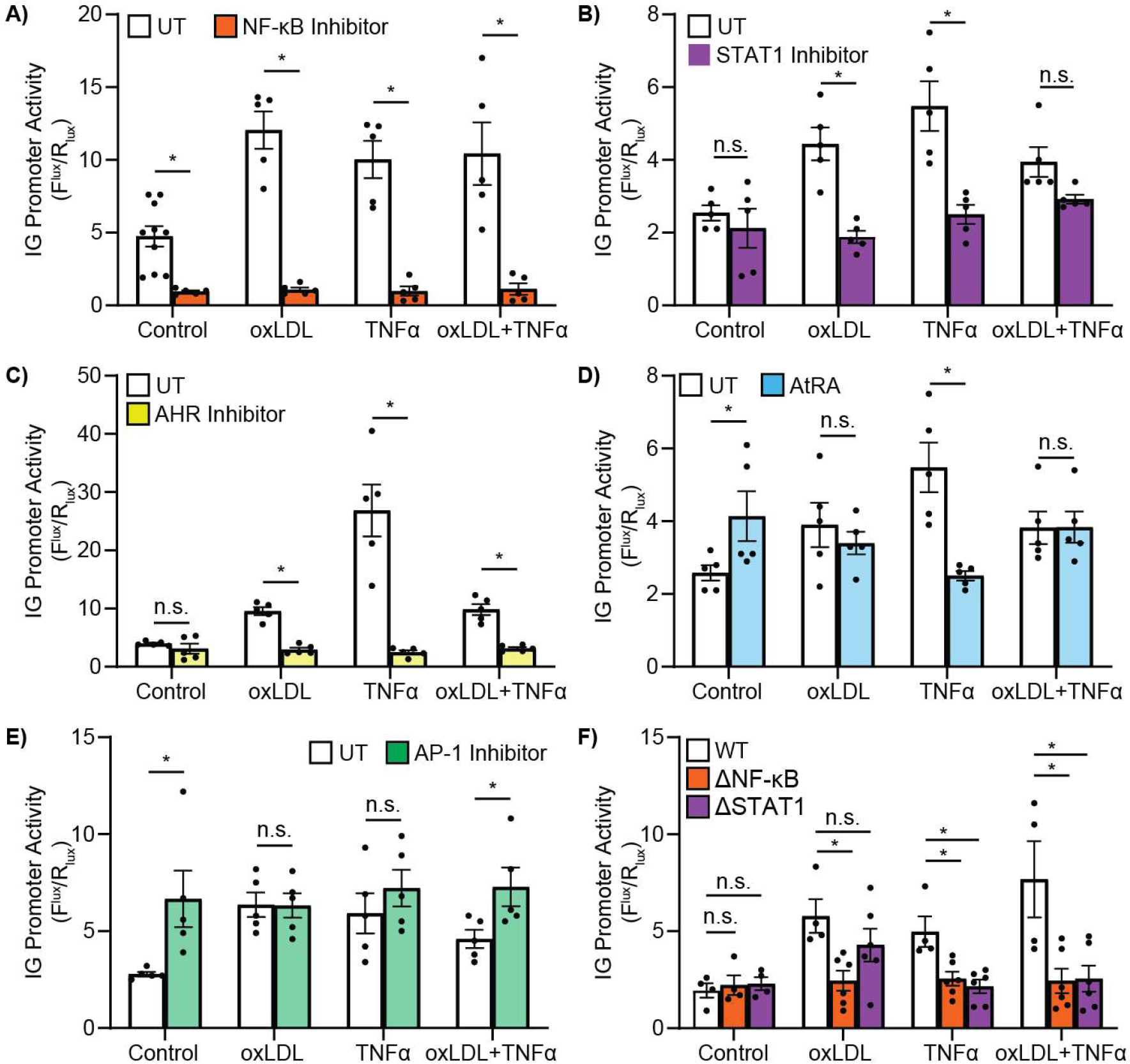
Transcriptional Regulation of the GATA2 IG Promoter in Macrophages by Atherogenic Stimuli. A dual-luciferase reporter of the GATA2 IG promoter was expressed in THP1-derived macrophages that were stimulated with 100 μg/mL oxLDL and/or 10 ng/mL TNFα for 48 hours. **A)** Impact of NF-kB inhibition with 20 μM of BMS-345541 on the activity of the IG promoter reporter. **B)** Impact of STAT1 inhibition with 10 μM nifuroxazide on the activity of the IG promoter reporter. **C)** Impact of AHR inhibition with 100 nM CH223191 on the activity of the IG promoter reporter. **D)** Impact of RORα activation with 1 μM all-trans retinoic acid (AtRA) on the activity of the IG promoter reporter. **E)** Impact of AP-1 inhibition with 1 μM SR11302 on the activity of the IG promoter reporter. **F)** Impact of deleting the putative NF-κB (ΔNF-κB, orange) or STAT1 (ΔSTAT1, magenta) binding sites from the wild-type (WT, green) dual-luciferase GATA2 IG promoter reporter. n = minimum of 5 biological replicates; * p < 0.05, n.s. = p > 0.05 between the indicated groups, ANOVA with Tukey correction.

### IDENTIFICATION OF GENES DIRECTLY REGULATED BY GATA2

Although we have previously characterized the impact of GATA2 overexpression on the transcriptome of macrophages, these experiments did not differentiate between genes regulated directly versus indirectly by GATA2^9,10^. Thus, we used THP-1 macrophages ectopically expressing GATA2 and ChIP-seq to identify the specific promoters bound by GATA2 in these cells. ChIP-seq data was analyzed with Galaxy Suite by first cleaning the data with trimmomatic and performing a quality check with FastQC of replicate samples^31,32^. Our ChIP-seq data consisted of ∼6 Gbp with sequence lengths from 20 to 101 bp (**Figure S1A**). Per base quality score was very good with an even distribution of all four nucleotides (**Figure S1B, C**). Fraction of reads in peaks was >1%, indicative of successful ChIP-seq (**Figure S1D**), and good agreement was observed across our duplicate samples (**Figure S2**)^33,34^. As expected, a strong GATA2 binding peak was found at the previously reported GATA2 binding site within the GATA2 promoter (**Figure S3A**), and MEME-ChIP analysis of the ChIP-seq dataset identified the expected GATA binding motif (**Figure S3B**)^35^. After confirming that the ChIP-seq met the ENCODE quality standards, peak calling was used to identify promoters bound by GATA2. >70% of precipitated genomic regions were associated with genes, with >50% of GATA2 binding sites located within 500 bp of a transcriptional start site (**Figure 3A, B**). Genes with binding peaks within 500 bp of the transcriptional start site—e.g. genes containing promoters bound by GATA2—were then analyzed with the Kyoto Encyclopedia of Genes and Genomes (KEGG)^36^. KEGG analysis found enrichment in genes for focal adhesion, cancer, MAPK signaling, and metabolic pathways (**Figure S3C-D**). We next determined the impact of GATA2 on the macrophage transcriptome by comparing this ChIP-seq data to our previously published transcriptomes of wild-type and GATA2-overexpressing THP-1 macrophages^9^. GATA2 bound to the promoters of 7,108 of the 20,424 transcripts expressed in these macrophages **(Figure 3D**). Of the GATA2-bound promoters, 10% were significantly upregulated (>2-fold change in expression), 7% were significantly decreased in expression, and the remainder exhibited less than a 2-fold change in their expression between wild-type and GATA2-overexpressing macrophages (**Figure 3D, E)**, with this analysis identifying SIGLEC6 as a marker of GATA2-expressing macrophages (**Figure S4**). Gene ontology analysis of the upregulated genes identified an enrichment in pathways for angiogenesis, proliferation, differentiation, chemotaxis, and the inhibition of apoptosis (**Figure 3F**), whereas the downregulated genes were enriched in pathways for responding to cytokines, lipid uptake and transport, foam cell development, and processes required for the removal of apoptotic cells (**Figure 3G**)^37^. Combined, these data indicate that GATA2 induces a transcriptional program that promotes the recruitment and proliferation of macrophages, while suppressing these cells sensitivity to cytokines and ability to engage in anti-atherogenic processes such as efferocytosis and lipid homeostasis.

**Figure 3:**
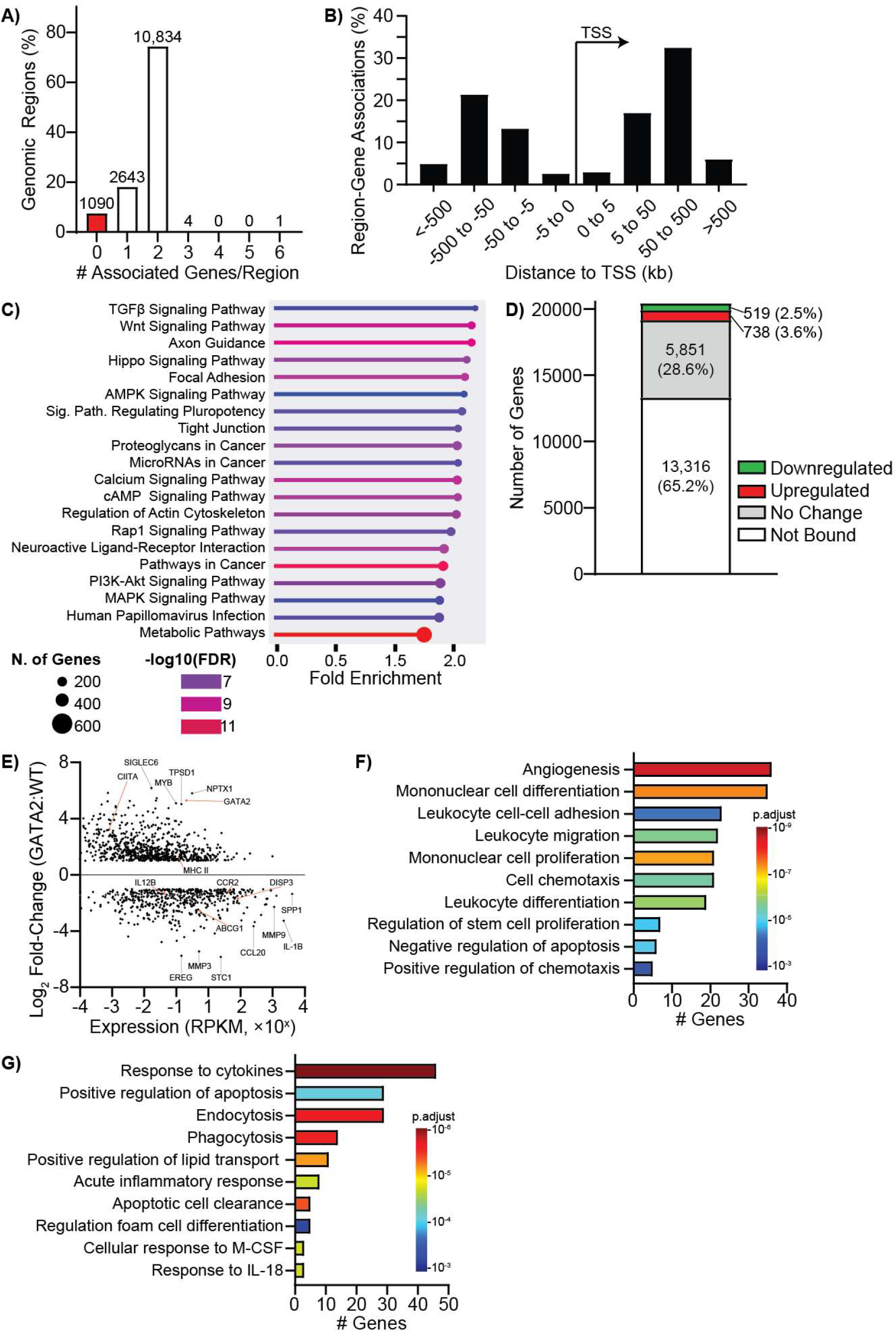
GATA2 ChIPseq Reveals Broad GATA2 Binding Across the Human Macrophage Genome. ChIP-seq was performed with GATA2-overexpressing THP1 derived macrophages, and the precipitated DNA analyzed to determine the pattern of GATA2 binding in these cells. **A)** Quantification of the number of GATA2 binding peaks that are associated with one or more genes. **B)** Distance from GATA2 binding peaks to the nearest transcriptional start site (TSS). Negative values indicate the binding site is located 5’ to the TSS; positive values indicate the binding site is located 3’ to the TSS. **C)** The biochemical function of genes near GATA2 binding peaks were classified using the Kyoto Encyclopedia of Genes and Genomes (KEGG) database to identify cellular pathways which are enriched for GATA2-bound genes. **D)** Comparison of the impact of GATA2 overexpression on the regulation of mRNAs expressed in GATA2-overexpressing versus wild-type THP1 macrophages versus the presence of a GATA2 binding peak at each gene. “Not Bound” indicates genes whose mRNA was detected in these cells, but to which GATA2 binding was not observed. “No Change” indicates genes where GATA2 binding proximal to the gene was detected, but the change in the abundance of the genes’ mRNA was less than 2-fold compared to wild-type cells. “Downregulated” and “Upregulated” indicate genes where there was a >2-fold change in mRNA abundance in GATA2-overexpressing versus wild-type macrophages. **E)** Comparison of the fold-change in gene expression between GATA2-bound genes in GATA2-overexpressing and wild-type THP1 macrophages compared to the expression level of those genes in wild-type cells. Genes showing the highest degree of differential expression are identified with black lines, genes identified as potentially impactful atherogenic genes are identified with orange lines. **F-G)** Gene ontology analysis of GATA2-bound genes which were upregulated > 2-fold (F) or downregulated >2-fold (G) in GATA2-overexpressing versus wild-type THP1 macrophages. Data summarizes the pooled data of 2 (ChIP-seq) or 3 (RNAseq) biological replicates.

### GATA2 REPRESSES MULTIPLE ANTI-ATHEROGENIC PATHWAYS

The KEGG and gene ontology data was further analyzed to identify genes associated with atherosclerosis or inflammation, identifying several genes associated with both processes. This included the cholesterol exporter ABCG1, the sterol-sensing inducer of lipid droplet formation DISP3, the shared p40 subunit of the inflammatory cytokine’s interleukin 12 and 23 (IL-12B), the chemokine receptor CCR2, major histocompatibility complex II (MHC II), and the transactivator of MHC II expression CIITA. Dualluciferase reporters containing the promoters of these genes were generated (**Table S1**) and expressed in either wild-type or GATA2-overexpressing THP-1 macrophages. All six of these genes showed promoter activity in wild-type THP-1 cells, with the DISP3 promoter showing the highest activity. Much to our surprise, GATA2 overexpression suppressed the promoter activity of all genes except CCR2 (**Figure 4A-E**). Moreover, GATA2 overexpression also negatively regulated the activity of the GATA2 IG promoter, indicative of a feedback look that restrains GATA2 activity (**Figure 4F**).

**Figure 4:**
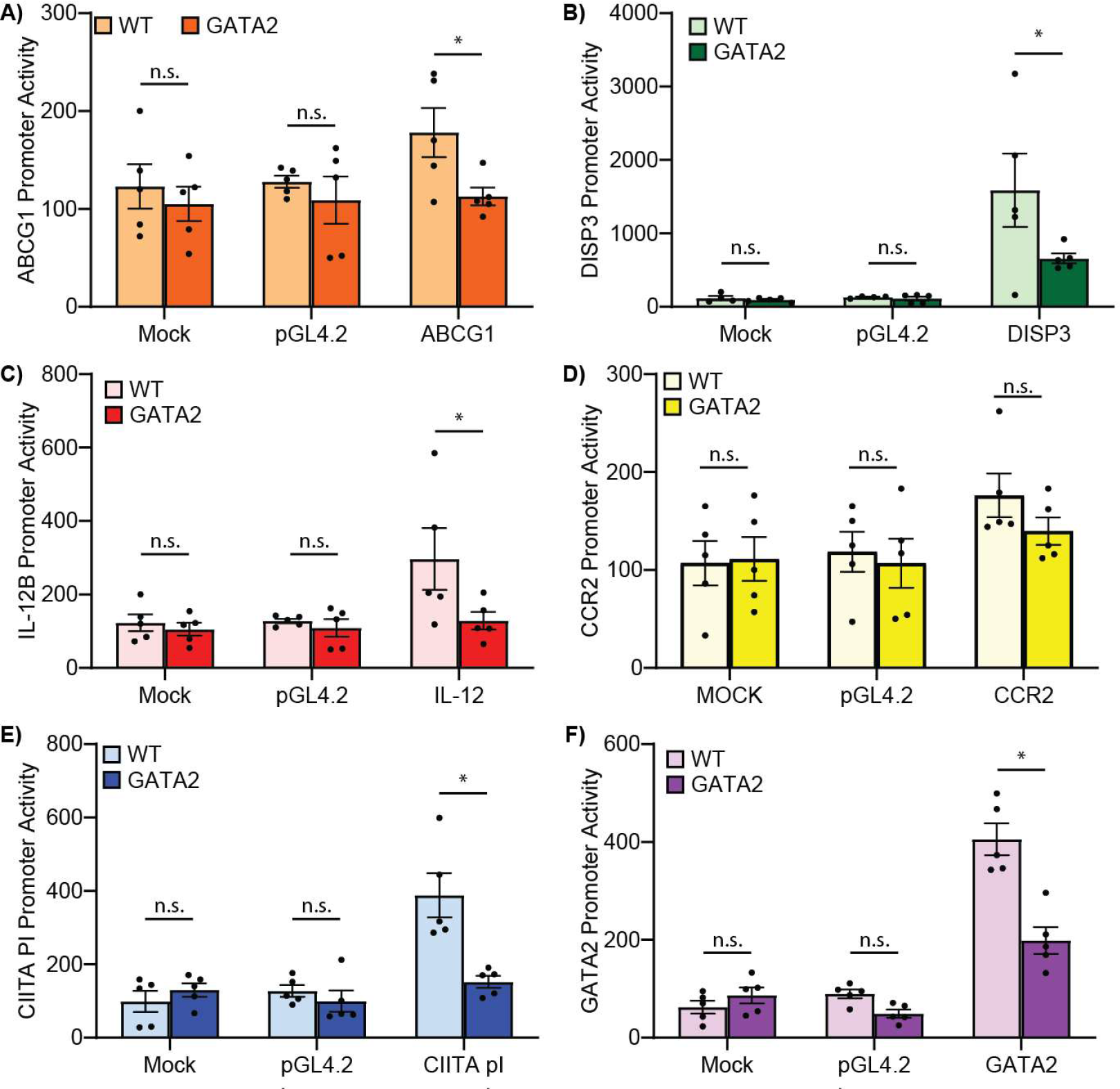
GATA2 Induces a Suppressive Transcriptional Landscape Across Several Atherogenic Loci. Dual-luciferase reporters were used to quantify promoter activity in wild-type (WT) versus GATA2-overexpressing THP1-derived macrophages. Luciferase activity is quantified in untrasnfected cells (Mock), cells transfected with a promotorless vector (pHL4.2), or transfected with luciferase vectors bearing the promoters for: **A)** the cholesterol exporter ABCG1, **B)** the lipid droplet and sterol-responsive protein DISP3, **C)** The common subunit of IL-12 and IL-23 (IL-12B), **D)** the chemokine receptor CCR2, **E)** the MHC II regulating transcription factor CIITA, and **F)** the GATA2 IG promoter. n = 5 biological replicates, * p < 0.05 between indicated groups, ANOVA with Tukey correction.

The promoter activity changes identified here suggests the presence of pervasive defects in macrophage function. Cholesterol homeostasis and efflux via ABCG1 was quantified using a cholesterol efflux assay, and consistent with the lower ABCG1 promoter activity in GATA2-overexpressing cells, GATA2 overexpression reduced cholesterol efflux by ∼20% (**Figure 5A**). Expression of IL-12B, the shared subunit of the inflammatory cytokines IL-12 and IL-23, was not affected by GATA2 over-expression (**Figure 5B**). Several receptors and signaling molecules required for the engulfment of pathogens (phagocytosis) and apoptotic cells (efferocytosis) were downregulated in GATA2 overexpressing macrophages, and consistently, both phagocytosis and efferocytosis were impaired by GATA2 expression (**Figure 5C**). Macrophages drive plaque expansion in part by generating reactive oxygen species (ROS) that then oxidize LDL and promote inflammation^38^. ROS production was assessed using the fluorogenic ROS sensor CellROX, comparing the basal production of ROS to ROS production after inflammatory (phagocytosis) and anti-inflammatory (efferocytosis) stimuli (**Figure 5D-E**). GATA2 overexpression significantly increased both basal and phagocytosis-induced ROS production by macrophages, but had no effect on the minimal ROS produced following efferocytosis (**Figure 5E**). CIITA is a transcription factor that is necessary for the expression of MHC II.

**Figure 5:**
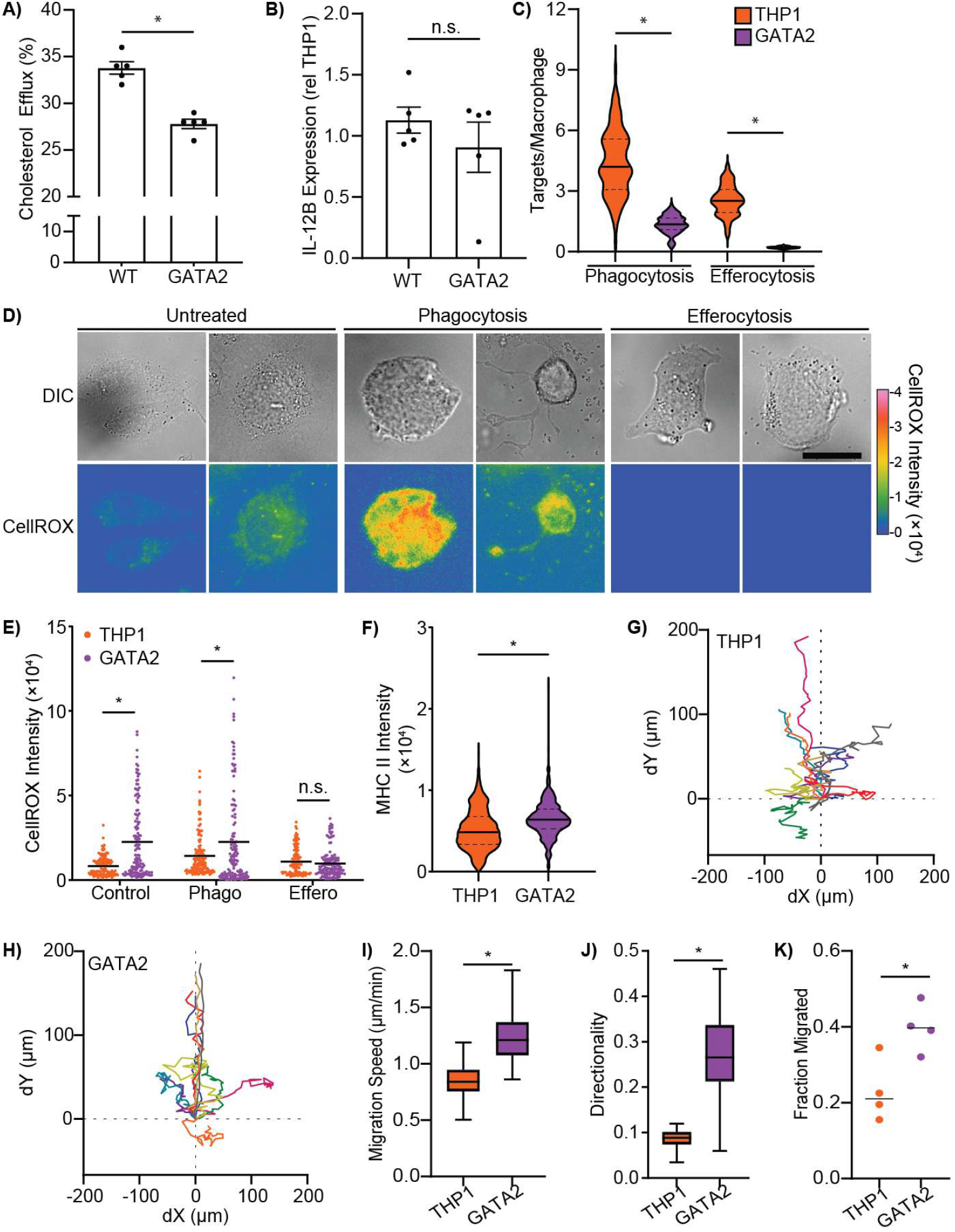
GATA2-Suppression of Gene Expression Impairs Macrophage Function. The activity of cellular pathways whose genes promoter activity were suppressed by GATA2 was assayed in wild-type (THP1) versus GATA2-overexpressing (GATA2) THP1-derived macrophages. **A)** Quantification of cholesterol efflux from macrophages to HDL over a 4-hour period. Efflux is quantified as the % of total cellular cholesterol that was exported from the cells. **B)** Expression of IL-12B, the common subunit of IL-12 and IL-23, by RT-qPCR. **C)** Phagocytosis and efferocytosis of particles mimicking antibody-opsonized pathogens (IgG coated beads, Phagocytosis) or mimicking an apoptotic cell (Phosphatidylserine-coated beads, Efferocytosis). Targets/cell for 120 cells/group, quantified across 3 biological replicates, are plotted. **D-E)** Representative micrographs (D) and quantification (E) of superoxide production in macrophages, as quantified by CellROX staining. Basal (untreated) superoxide production, as well as superoxide production induced 1 hour after the phagocytosis of *E. coli* or efferocytosis of apoptotic cells is measured. Each dot quantifies the integrated CellROX signal from a single macrophage. 75 cells, imaged across 3 biological replicates, are quantified in each group. **F)** Single-cell quantification of MHC II expression in macrophages, >600 cells/group were quantified in 3 biological replicates. **G-K)** Impact of GATA2 expression on THP1 macrophage chemotaxis to 50 ng/mL CCL2 in a microfluidic chemotaxis chamber, showing 10 representative migratory tracts from wild-type (G) and GATA2-overexpressing (H) macrophages, with the speed (I), directionality (J), and portion of macrophages capable of responding to CCL2 (K) quantified from >200 cells/group, imaged across 3 biological replicates. n = 5 (A,B) or 3 (C-K) biological replicates. * = p < 0.05 between indicated groups, Students *t-*test (A, B, F, I-K) or ANOVA with Tukey correction (C,E).

## Discussion

GATA2 is upregulated in macrophages within atheromas, with mutations in GATA2 predisposing patients to atherosclerosis via an unidentified mechanism^9,12,35^. In this study we investigated the regulation of GATA2 and the impact of GATA2 expression on the transcriptome and function of macrophages, using ectopic expression of GATA2 to avoid the confounds of Unexpectedly, we observed a >50% decrease in the promoter activity of the CIITA pI (macrophage-specific) promoter (**Figure 4E**), but we observed an ∼30% increase in the cell-surface density of MHC II (**Figure 5F**). Lastly, using a microfluidic chemotaxis chamber we quantified the migration of macrophages to the chemokine CCL2. GATA2 overexpression significantly enhanced chemotaxis, increasing the speed and directionality of migration, and the portion of cells which responded to the chemokine (**Figure 5G-K**). Combined, these data indicate that GATA2 overexpression generates a pro-atherogenic phenotype in macrophages through mediating a loss of cholesterol homeostasis, enhancing inflammatory and oxidative pathways, and driving an invasive migratory phenotype.

GATA2-inducing stimuli such as oxLDL. GATA2 expression induced a stem cell-like phenotype in macrophages by mediating the upregulation of genes associated with proliferation and differentiation. Functional analyses of GATA2 expressing macrophages revealed several cellular defects which would contribute to the development of atherosclerosis, including reduced cholesterol efflux capacity, elevated superoxide production, and a more invasive/chemotactic phenotype, while our previous studies demonstrated that GATA2 expression promoted macrophage proliferation and suppressed efferocytosis^9,10^. Combined, these data identify GATA2 as a driver of several transcriptional and cellular pro-atherogenic pathways mediated by GATA2 expression in macrophages.

Macrophages contribute to atherosclerosis through a variety of mechanisms. Most of the macrophages in the plaque are derived from recruited monocytes which proliferate and differentiate into macrophages, with local proliferation producing the majority of plaque macrophages^6,39^. These macrophages can take on several atherogenic and atheroprotective polarization states, each driven by a unique transcriptional program. ATF-1 expression, induced in plaque macrophages by the presence of free heme, mediates the polarization of macrophages to an Mhem phenotype^8^. These atheroprotective macrophages then stabilize the plaque by removing the oxidation-promoting heme and enhancing cholesterol efflux out of the plaque. In contrast, ATF2 activity drives the polarization of macrophages into atherogenic foam cells which accumulate lipids and secrete proinflammatory cytokines that further drive plaque inflammation and growth^40^. Unlike these macrophage subtypes, we did not observe a change towards a distinct pro- or anti-atherosclerotic phenotype in the GATA2+ expressing macrophage population. While some atherogenic changes were observed – e.g. impairments in lipid transport – these cells also exhibited changes consistent with an atheroprotective phenotype such as decreased sensitivity to cytokines and decreased expression of genes involved in foam cell development. Our data indicates that GATA2 directly upregulates genes involved in proliferation, differentiation, and in the inhibition of apoptosis, and indeed, we have previously observed the enhanced proliferation and resistance to apoptosis of GATA2-expressing macrophages^10^. Similarly, many macrophage-specific processes are downregulated by GATA2 including phagocytosis, cholesterol export, and inflammatory pathways. This biasing of macrophages towards proliferation, and downregulation of genes that regulate prototypical macrophage activities such as phagocytosis, suggests that GATA2 is inducing a stem cell-like phenotype in these cells, perhaps in a manner similar to the macrophage-to-monocyte dedifferentiation observed in *Bordetella pertussis* infection^41^. This is consistent with the role of GATA2 in the hematopoietic and lymphatic vessel compartments. In both of these compartments, GATA2 expression promotes a stem cell-like phenotype typified by proliferation and a block on cells entering terminal differentiation pathways^14,42,43^. Indeed, mutations which lower GATA2 expression led to a loss of monocytes, macrophages, and other myeloid cells due to a collapse of the myelopoietic compartment, with GATA2 deficiency resulting in a total loss of the hematopoietic stem cell population and a collapse of hematopoiesis^44,45^. Similarly, reduced GATA2 expression impairs the growth and regeneration of lymphatic vessels, where GATA2 is required to mediate both the differentiation of progenitors into lymphatic endothelial cells, and the proliferation of lymphatic endothelial cells^43,46^. Indeed, our ChIP-seq and RNAseq data identified multiple genes bound and upregulated by GATA2 that are associated with stemness and self-renewal of hematopoietic stem cells including RUNX2, SHH, PBX1, and TERT^47–50^. Consistent with an immature phenotype, we observed an upregulation in differentiation and proliferation genes in GATA2-overexpressing cells, consistent with incompletely developed myeloid cells.

Immature myeloid cells are frequently found in chronically inflamed sites, including in the synovium of rheumatoid arthritis patients, within tumors, and in atherosclerosis^51–54^. While it is unclear why seemingly immature cells are present at these sites, it has been suggested that the stress of on-going inflammation results in “emergency myelopoiesis” that prematurely releases developing myeloid cells into the circulation^52^. When recruited into inflamed sites, these immature cells promote further inflammation through processes such as inhibiting the function of Treg cells, producing reactive oxygen species, promoting granulopoiesis, recruiting Th17 T cells, and producing inflammatory cytokines such as IFNγ and IL-12^51,54–57^. However, not all immature myeloid cells are inflammatory, with myeloid-derived suppressor cells (MDSCs)—also derived from immature myeloid cells—arising under the same chronic inflammatory conditions, but which serve to suppress inflammation. While MDSCs may start as immature myeloid cells, establishment of the anti-inflammatory MDSC phenotype requires pathological activation in peripheral tissues^58,59^. Transcriptomic analysis of MDSC differentiation did not identify GATA2 expression, suggesting that the GATA2+ macrophages we have identified are not MDSC cells, and indeed, our cells lacked expression of the MDSC markers CD84 and VEGFA^60–62^. Moreover, the stem-like phenotype, upregulation of proliferative and cell-survival pathways, and the induction of this phenotype by oxLDL and TNFα, suggest that these cells are taking on a stem-like phenotype within the plaque environment. While this does not exclude the possibility these cells arose from either immature myeloid cells or MDSCs, their stem-like gene expression pattern, suppressed production of IL-12, and their lack of expression of MDSC markers, suggests that these GATA2+ macrophages represent a unique population of stem-like macrophages that deviate significantly from both immature (inflammatory) myeloid cells and from MDSCs.

While GATA2 is known to play important roles in development, hematopoiesis, and the maintenance of lymphatics, the specific genes regulated by GATA2, and the biological impact of this regulation, has been difficult to uncover. Indeed, only a handful of genes have been definitively shown to be regulated by GATA2, including the androgen receptor, FAR2, GATA1, HDC, c-KIT, and GATA2 itself ^63–65^. In contrast, several genes have been shown to be downregulated by GATA2, although in many cases this appears to occur independently of GATA2 binding to these genes or their promoters^65^. This difficulty stems from the manner in which GATA2 functions. As a pioneering transcription factor, GATA2 can open chromatin to enable access by other transcription factors – but GATA2 may have no further discernable impact on those gene(s) expression as other transcription factors then regulate expression in the now-open chromatin^31,66^. This phenomenon may explain why we observed GATA2 binding to over 5,800 loci across the genome, but with only 20% of these loci showing differences in gene expression. Moreover, GATA2 often acts in concert with other transcription factors such as GATA1, CEBPA, SPI1, and ZFPM1, and consequently some of the loci that we found were bound by GATA2 may not exhibit changes in expression due to the absence of one of these partners^67^. As an example, we observed decreased expression of CIITA, but increased expression of its target gene MHC II. This may be a product of GATA2 inducing expression of CIITA from one of its two non-myeloid promoters (pIII or pIV), although these promoters appear to lack GATA2 binding sites, meaning that this interaction is likely indirect^68–70^. An additional complicating factor is that GATA2’s activity is concentration dependent, with higher GATA2 concentrations promoting the formation of a GATA2 homodimer which has a different activity than the GATA2 monomer^71^. Moreover, GATA2’s ability to occupy a locus is also often concentration-dependent due to competition for binding by other GATA family transcription factors. This competition results in many promoters being regulated by a “GATA switch”, wherein different GATA transcription factors compete for binding to the same G-A-T-A DNA motifs, with occupancy of the motif determined by the relative concentration of the competing GATA transcription factors^63^. Indeed, this GATA-switch is central to hematopoiesis, GATA2 is displaced from multiple loci in progenitor cells by the increasing expression of GATA1, allowing the progenitor to cease self-renewal and enter terminal differentiation^72–74^. Further complicating our understanding of GATA2’s impact on the genome is the large number of microRNAs whose expression is dependent on GATA2^66,75–77^. This allows GATA2 to regulate a broad array of genes without interacting with those genes’ promoters. As one example, GATA2 upregulates mIR126 in lymphatic endothelium, in turn downregulating claudin 5 and VE-cadherin thus causing lymphatic junctional defects^78^. This complex biology likely contributes to some of the discrepancies we observed between GATA2’s impact on a gene’s promoter activity versus the mRNA levels of those genes observed in the same cells.

Our data indicate that GATA2 expression impacts a large portion of the macrophage transcriptome, leading to a stem-like phenotype typified by increased macrophage proliferation and survival. This stem-like state results in the loss of core macrophage functions including cholesterol export, phagocytosis, and efferocytosis, while simultaneously effecting a modest proinflammatory state through increased ROS production and cytokine secretion. The net effect of these changes is a pro-atherogenic phenotype, with these cells exhibiting loss of cholesterol export and efferocytic capacity that is required for disease onset and progression, while also promoting a proliferative phenotype which contributes to the cellularity of atherosclerotic plaque. As such, GATA2 may be a suitable target for the development of future atherosclerosis therapies.

## Materials & Methods

### Materials

pGL4.20[luc2/Puro] was a gift from Dr. Rodney DeKoter, University of Western Ontario. Lympholyte poly and THP-1 human monocytic cells (ATCC TIB 202) were purchased from Cedarlane Labs (Burlington, Canada). THP-1 cells transduced with a GATA2 overexpression cassette were generated previously ^9^. Information on all antibodies used in this study can be found in **Table S2**. Information on all primers used in this study can be found in **Table S3**. Primers and synthetic DNA fragments were from IDT (Coralville IA). DMEM, RPMI, 100× antibiotic/antimycotic, and fetal bovine serum (FBS) were purchased from Wisent (Saint-Jean-Baptiste, Canada). Cholesterol efflux kit and BMS-345541 were from Abcam (Cambridge, UK). 96-well luciferase plates were from Greiner Bio-One (Kremsmünster, Austria). SR11302, all-trans-retinoic acid, CH223191, and nifuroxazide were from Cayman Chemical (Ann Arbor, MI). Coverslips, 16% paraformaldehyde (PFA), and immersion oil were from Electron Microscopy Sciences (Hatfield, PA). CCL1, CellROX, oxLDL, formaldehyde, and TNFα were purchased from ThermoFisher (Waltham, MA). Amaxa nucleofector kits were purchased from Lonza Canada (Kingston, Canada). Luciferase detection kits were from Promega (Madison WI). RNA isolation kits were from Qiagen (Hilden, Germany). iScript cDNA kits and SsoFast Evagreen Supermix were from Biorad (Hercules, CA). Megna-ChIP kit and HALT protease inhibitors were from Millipore-Sigma (Burlington, Ma). µ-Slide Chemotaxis chambers were from Ibidi (Gräfelfing, Germany). SMART RACE 5’/3’ and NucleoSpin Gel and PCR Clean-Up kits were from Takara Bio (Ann Arbor, MI). RNAeasy Mini Kit was from Qiagen (Toronto, Canada). Prism software was from Graphpad Software (Boston, MA). All other common laboratory chemicals were purchased from Bioshop Canada (Mississauga, Canada).

### THP-1 Cell Culture

THP-1 human monocytes, both wild-type and cells ectopically over-expressing GATA2, were maintained as suspension cultures in RPMI + 10% FBS in a 37°C/5% CO_2_ incubator. Cells were grown to a density of 1 × 10^6^/mL and split by diluting 1:5 into fresh medium. THP-1-derived macrophages were differentiated from by first placing 2.5 × 10^5^ cells into each well of a 12-well plate, and incubated in RPMI + 10% FBS + 100 ng/mL PMA for 72 hr. For experiments involving microscopy, #1.5 thickness, 18 mm diameter circular coverslips were cleaned by acid washing (2 M HCl in ddH_2_O, 56°C, overnight), autoclaved, and then placed into the wells of the 12-well plate prior to plating cells.

### Macrophage Transduction and Stimulation

THP-1 cells were transfected with 0.5 μg of DNA using an Amaxa Human Monocyte Nucleofector kit and a Lonza Nucleofector II, using the manufacturer’s V-001 protocol. After transfection, 500 µL of warm RPMI + 10% FBS was added to the electroporation cuvette. Transfected cells were then transferred to a 12-well plate and incubated in a humidified 37^°^C/5% CO_2_ incubator overnight, and macrophage differentiation induced as described above. After 72 hrs, media was removed from wells and fresh RPMI + 10% FBS media was added to macrophages with oxLDL (100 μg/mL) or TNFα (10 ng/mL) or a mixture of oxLDL + TNFα for 48 hrs. For inhibition studies, inhibitors of NF-kB (BMS-345541, 20 μM), AP-1 (SR11302, 1 μM), STAT1 (nifuroxazide, 10 μM), or AHR (CH223191, 100 nM) were added at the same time as oxLDL/TNFα. ROR activation was induced by the addition of all-transretinoic acid (1 μM) at the time of oxLDL/TNFα addition. After stimulation and/or inhibition, macrophages were cultured for an additional 48 hrs in a humidified 37°C + 5% CO_2_ incubator.

### RNA Isolation, 5’ RACE, AND RT-QPCR

RNA was extracted from THP-1 macrophages using a RNeasy Mini Kit according to manufacturer protocol and RNA integrity confirmed by Bioanalyzer (Agilent Technologies). Only RNA with an RNA integrity number of 7 or higher was used for subsequent experiments. cDNA was synthesized using a SMARTScribe Reverse Transcriptase kit for 5’ RACE, or with an iScript kit for RT-qPCR. 5’ RACE was performed using a SMART RACE 5’/3’ kit as per manufacture’s instructions. Briefly, a GATA2-specific primer was generated that bound to the first coding exon of GATA2 was generated (**Table S3**) and added to the cDNA along with universal primer mix A and SeqAMP DNA polymerase, and run through 35 PCR cycles with denaturation at 94°C/30 sec, annealing at 68°C/30 sec, and elongation of 72°C/3min. The resulting 5’ RACE product was resolved on a 1.2% agarose gel. The 5’ RACE product was purified using a NucleoSpin Gel and PCR Clean-Up Kit as per the manufacture’s protocol and sequenced by Sanger sequencing at the London Regional Genomics Facility (London, Canada) and aligned to the human hg38 genome database using BLAT tool within the UCSC genome browser. For RT-PCR, SsoFast Evagreen Supermix with THP-1 cDNA used as template along with gene-specific primers (**Table S3**) as per manufacturer’s instructions, with amplification and analysis performed using a QuanStudio qPCR machine (ThermoFisher).

### Chromatin Immunoprecipitation Sequencing and Analysis

ChIP-seq was performed using a Megna-ChIP kit as per the manufacture’s instructions. 2.0×10^7^ differentiated GATA2-overexpressing THP-1 cells were cultured in a 150 mm culture dish, washed three times with PBS, and fixed for 10 min with 1% formaldehyde. 1% glycine was added to quench unreacted formaldehyde, then the cells washed with ice-cold PBS. The cells were then scrapped into cold PBS with HALT protease inhibitors, centrifuged for 5 min at 300 × g, and the cells resuspended in lysis buffer and incubated on ice for 15 min with mixing every 5 min. The lysate was cleared with a brief centrifugation, and suspended in nuclear lysis buffer. DNA was then sheared by sonicated with a Bioruptor (Diagenode, Denvile NJ), using 15 cycles of a 30 seconds on/30 seconds off to generate fragments with the desired ∼300 bp length (data not shown), then the samples centrifuged to remove insoluble materials. Rabbit anti-human GATA2 antibody (**Table S2**) and 20 μL of protein A/G magnetic beads were added 50 μg of sheared DNA and incubated overnight at 4°C on a rotating shaker. Protein A/G beads were separated using a magnetic separator and the supernatant removed. The protein A/G beads were then sequentially washed with low salt wash buffer, high salt wash buffer, LiCl wash buffer, and TE buffer, using the magnetic separator after each wash to remove the beads. Elution buffer with proteinase K was then added and incubated at 62°C for 2 hrs, then incubated at 95°C for 10 min. The beads were removed via magnetic separation, and the DNA purified from the supernatant using the Megna-ChIP kit. DNA concentration was quantified using Qubit (ThermoFisher Scientific, Mississauga, Canada). Library generation and sequencing of samples and input were performed at the Montreal Clinical Research Institute genomics facility using 15 ng of recovered DNA using the KAPA HyperPrep kit according to the manufacturer’s protocol. The quality of libraries was checked, and libraries were sequenced on a PE100-50 M reads run on a NovaSeq 6000 (Illumina, San Diego, CA) and the data exported in FASTQ format. Raw data (FASTQ files) are available from Borealis, dataset HW5AHK, https://doi.org/10.5683/SP3/HW5AHK.

ChIP-seq data was analyzed using Galaxy Suite^31^. Quality control was performed using FastQC, then cleaned and adaptor sequences removed using Trimmomatic. Peak calling was performed on the trimmed data using MACS2 callpeak, using input DNA (prior to GATA2 immunoprecipitation) as a control. For peak detection, NarrowPeaks was called using the *H. sapiens* (2.7 × 10^9^) genome size with single-end BAM file, and a shifting model built using a lower mfold bound of 5, upper mfold bound of 50, bandwidth fragment size of 300, and minimum false discovery rate cutoff of 0.05. Peaks were viewed using a Bigwig file that was generated with GRch38/hg38(2701495761) using bamCoverage. GREAT was used to get peaks to genes association using Human:GRCh38 (UCSC hg38, Dec 2013) assembly and basal plus extension setting (5.0kb upstream, 1.0kb downstream, and 1000.0kb maximum extension). The quality of peaks was checked using the ChIPQC package on R Studio^79^. FASTA DNA was extracted from peak calling tabular form using Extract Genomic DNA. MEME-ChIP was used for motif analysis and ShinyGO was used for pathway analysis with an FDR cutoff of 0.05.

To compare this ChIP-seq data to our previously published RNAseq data of the same wild-type and GATA2-derived macrophages, the list of genes bound by GATA2 were exported using the standard Human (GRCh38/hg38) gene nomenclature, and imported alongside the RNAseq data into Matlab. Genes appearing in both databases were identified by their GRCh38/hg38 designation, and the mean RPKM for wild-type and GATA2-overexpressing macrophages calculated. Genes were then classified as unbound (not bound by GATA2 but appearing in the RNAseq dataset), no change (genes bound by GATA2 in the ChIP-seq dataset but exhibiting a <2-fold difference in gene expression in the RNAseq dataset), or down- or up-regulated (>±2 fold change in expression).

### Luciferase Assays

Genes whose promoters were bound by GATA2 were identified from the ChIP-seq dataset and prioritized to identify genes with known roles in atherosclerosis, inflammation or lipid processing. Luciferase vectors for the pI (macrophage-specific) promoter for CIITA was generated previously^70^. The promoters of high-priority genes were identified in the Human GRCh38/hg38 genome assembly using the UCSC genome browser, and the promoter regions identified as the regions surrounding the transcriptional start site containing elevated levels of Histone H3-Lysine 27 acylation, a histone mark indicative of active promoters and enhancer regions^80^. The sequence of these regions were imported into SnapGene software (**Table S1**), and regions of homology with the 20 bp of the pGL4.20[luc2/Puro] on either side of the EcoRV site added to both ends of the promoter sequences. These sequences were then ordered as synthetic DNA constructs and inserted into EcoRV-digested pGL4.20[luc2/Puro] vectors via Gibson assembly using a HiFi Gibson Assembly Kit as per the manufacture’s instructions. The resulting vectors were sequenced using Sanger sequencing at the London Genomics Centre (London, Canada) to confirm successful assembly. THP-1 cells were transfected with 0.5 μg of DNA and differentiated to macrophages as described above. Following differentiation, the cells were lysed with 250 µL passive lysis buffer followed by gentle mixing for 20 mins at room temperature. Lysates were collected, cleared with a 21,000 × g/30 sec centrifugation, and 100 µL of the supernatant added per well of a 96-well white polystyrene plate. To each well 50 µL of luciferase assay reagent II was then added, and the plate immediately read using 2(L) Lum, starting with 10 sec linear shaking in Cytation Gen5 software. Duplicate wells for all measurements were used, with each biological replicate being the average of these duplicate technical repeats.

### RT-PCR

Total RNA was isolated from THP-1 cells using a Qiagen RNA isolation kit and cDNA generated with an iScript cDNA kit. q-PCR was performed using SsoFast Evagreen Supermix on a Quantico Q PCR thermocycler. Gene expression was calculated using the ΔΔCt method, using GAPDH as a reference gene. All q-PCR primer sequences are in **Table S3**.

### Immunofluorescence

THP-1 macrophages were differentiated in the wells of 12-well plates into which 18 mm diameter, #1.5 thickness round coverslips had been placed. For CellROX staining, 5 μM of CellROX Deep Red was added to each well and incubated at 37°C and 5% CO_2_ for 30 min. The culture medium was then removed and the cells fixed by addition of 4% paraformaldehyde (PFA) in PBS for 20 min at room temperature. The PFA was then removed, the cells washed with PBS, and the coverslips mounted on slides using Permafluor mounting media prior to imaging. For immunostaining, the culture medium was removed from differentiated macrophages and the cells fixed by with 4% paraformaldehyde as described above. After fixation the cells were washed with PBS and then blocked with antibody buffer (2% bovine serum albumin in PBS) for 1 hr. Primary antibodies were added in antibody buffer at the concentrations indicated in **Table S2** and incubated for 1 hr at room temperature with gentle rocking. The cells were then washed 3 × 15 min in PBS at room temperature with gentle rocking, followed by addition of a 1:1,000 dilution of labeled secondary antibodies for 1 hr at room temperature. The cells were then washed 3 × 15 min in PBS. Some samples were then counterstained by addition of 5 μg/mL Hoechst 33342 in PBS for 5 min. After staining the coverslips were mounted on slides using Permafluor mounting media. All samples were imaged on a Zeiss AxioObserver microscope equipped with 100×/1.40 NA and 63×/1.40 NA objectives, a colibri excitation light source, Hamamatsu ORCA-Fusion CMOS camera, DIC white light illumination, and operated using Zen 3.12 software. Identical excitation intensity and exposure times were used to allow for intensity comparisons between samples labeled in the same biological replicate. Tiled images containing at least 200 cells were captured of each sample, and the images exported as OME-formatted TIFs. Individual cells were identified in each image using a trained pixel classification algorithm in Ilastik and the resulting segmented mask exported^81,82^. The segment mask was then loaded in FIJI and the “Analyze Particles” feature used to generate ROIs for all cells in the image^83^. These ROIs were then used to quantify the integrated intensity of the fluorescent label in each cell.

### Cholesterol Efflux Assay

Cholesterol efflux was quantified using a Cell Based Cholesterol Efflux Assay Kit as per the manufacture’s instructions. Briefly, THP-1 macrophages were differentiated at 90% confluency in the wells of a 96-well plate, using duplicate wells for each condition. Once differentiated, the culture medium was removed from each well and replaced with 50 μL of labeling medium mixed with an equal volume of serum-free RPMI, and the cells incubated for 1 hr at 37°C in a 5% CO_2_ humidified incubator. The labeling medium was replaced with 100 μL of equilibrium medium and incubated for 16 hrs in a 37°C/ 5% CO_2_ humidified incubator. The cells were then washed with 200 μL of phenol-red free/serum-free RPMI, and 100 μL of phenol-red free/serum-free RPMI medium containing 2% (v/v) LDL/VLDL-depleted human serum and the cells incubated for 4 hrs in a 37°C/ 5% CO_2_ humidified incubator. The supernatant was then transferred to a white 96-well plate, the cells lysed with the included lysis buffer, and the cell lysate transferred to separate wells in the white 96-well plate and fluorescence measured on a fluorescent plate reader using 485 nm excitation and 523 nm ± 5 nm emission. Cholesterol efflux was calculated as the fluorescence intensity of the supernatant divided by the sum of the fluorescence intensity of the supernatant and cell lysate.

### Phagocytosis and Efferocytosis Assays

Phagocytosis and efferocytosis were quantified as per our published protocols^84,85^. Briefly, phagocytic targets were generated by washing 10 μL of 3 μM diameter DVB-PS beads 3 × 1 min PBS, using 5000 × g centrifugation to collect the beads after each wash. Washed beads were then incubated for 1 hr at room temperature in 100 μL PBS + 10 μg of purified Rat IgG. The beads were washed 3× with PBS and diluted into 100 μL of PBS. Efferocytic targets were generated by washing 3 μM dimeter silica beads 3 × 1 min PBS, using 5000 × g centrifugation to collect the beads after each wash. At the same time 0.04 mmol of a 20:80 molar ratio of phosphatidylserine:phosphatidylcholine was prepared and dried under a flow of nitrogen. The washed beads were then suspended in 1 mL of PBS, this suspension used to resuspend the dried lipid mixture, and the combined mixture incubated for 1 hr at room temperature. The beads were washed 3× with PBS and diluted into 100 μL of PBS. 2.5 × 10^5^ THP1-derived macrophages were differentiated on coverslips as described above, and 3 μL of the bead suspension (∼20:1 bead:macrophage) was placed into each well. The cells were then incubated for 60 min in a 37°C/5% CO_2_ incubator, then washed once with PBS and fixed with 4% PFA in PBS at 37°C for 15 min. The cells were then rinsed 3 × with PBS, and a 1:750 dilution of anti-Rat IgG-Cy3 (IgG coated beads) or AnnexinV-AlexaFluor555 in imaging buffer (150 mM NaCl, 5 mM KCl, 1 mM MgCl_2_, 100 μM EGTA, 2 mM CaCl_2_, 20 mM NaHCO_3_, pH 7.4) and immediately imaged. Beads were identified using the DIC channel, and non-internalized beads identified via Cy3/AlexaFluor555 staining, and the number of fully internalized beads in each macrophage quantified.

### Superoxide Measurements

THP-1-derived macrophages were cultured on coverslips in a 12-well plate as described above. Phagocytic targets were generated by culturing RFP-expressing BL21 *E. coli* overnight in LB medium + 50 μg/mL kanamycin. The culture was diluted 1:10 in fresh LB, the OD_600_ measured, 1 × 10^8^ cells collected and washed with PBS and fixed with 4% PFA for 30 min. The bacteria were then washed twice with PBS and resuspended in PBS at 1 × 10^8^ cells/mL. Efferocytic targets were prepared as above. For each well, 3 µL of efferocytic targets or 7.5 × 10^6^ bacterial cells were added to 1 mL of pre-warmed RPMI + 10% FBS. Cells were washed three times with PBS before adding the prepared media with beads or bacteria. The plate was centrifuged at 100 × 6 for 1 min and incubated at 37°C for 10 min. The cells were then gently washed three more times with PBS, followed by the addition of 1 mL of fresh, warmed media per well. The plate was then incubated at 37°C for 1 hour. After incubation, the media was removed, 5 µM CellROX Deep Red added, and the cells incubated for an additional 30 min at 37°C. The cells were then washed 3× with PBS and fixed with 4% PFA for 15 minutes. After an additional three PBS washes, cells were mounted on coverslips with Permafluor and the samples imaged using a 63×/1.40NA objective on a Zeiss Axiobserver microscope, and the resulting images analyzed with FIJI.

### Chemotaxis Assays

2.5 × 10^5^ wild-type or GATA2-overexpressing THP-1 were differentiated into macrophages in a 12-well plate as described above. After differentiation, cells were removed from the plate by washing once with PBS pre-warmed to 37°C, and then 300 μL of Versine applied to each well. After a 5 min/37°C incubation, the cells were gently scraped with a cell scraper, 700 μL of complete medium added, the cells transferred to a 1.5 mL microcentrifuge tube, and the cells pelleted with a 1,000 × g/ 3 min centrifugation. The cell pellet was then suspended in 10 μL of complete medium and transferred into the central channel of an Ibidi µ-Slide Chemotaxis chamber^86^. The chamber was then placed in a petri dish next to two whetted kimwipes and incubated in a 37°C/5 % CO_2_ incubator for 1 hr. The central channel was then washed 2× with 10 μL serum-free RPMI to remove non-adherent cells, then both reservoirs filled with 65 μL serum-free RPMI and the chamber incubated an additional 2 hrs in a 37°C/5 % CO_2_ incubator. To one reservoirs, 30 μL of RPMI + 100 ng/mL of recombinant human CCL2 was added, then the chemotaxis chamber placed on the heated and CO_2_-perfusd stage of our Zeiss AxioObserver microscope. Cell migration was then imaging using DIC images of the central channels were then recorded with DIC illumination and a 10×/0.30 NA objective lens, taking 1 image every 5 min over 18 hr. The resulting images were exported, cells segmented using a trained algorithm in Ilastik, and the migration of cells tracked using the TrackMate package in ImageJ^87,88^. Two measurements of chemotaxis were made, on an individual cell basis:

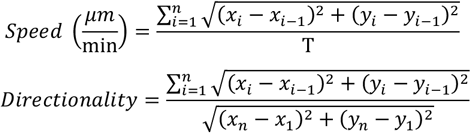

where x_i_/y_i_ = x/y position of the cell at timepoint I, and ⊤ is the duration of the experiment.

### Statistics

Statistics were performed in Graphpad Prism version 9. Data is presented as individual independent biological repeats plotted over the mean ± SEM. All data is tested for normality using a Shapiro-Wilk test prior to analysis. Unless otherwise noted, all statistics were performed using non-parametric 2-tailed test, and with a significance cutoff of 0.05. The specific statistical tests used are indicated in the figure captions.

## Acknowledgements & Funding

We would like to thank the Molecular Imaging Facility (www.molimage.ca) and Kristin Chadwick of the London Regional Flow Cytometry Facility for their assistance with the microscopy and flow cytometry assays. This study was funded by a Heart and Stroke Foundation of Canada Grant-In-Aid to BH. AA was funded by an Ontario Graduate Scholarship and a Dr Robert George Everitt Murray Graduate Scholarship from The University of Western Ontario. PHC was funded via the X-Labs program at The University of Western Ontario. The funders had no role in study design, data collection and analysis, decision to publish, or preparation of the manuscript.

