## Supplemental Figures and Tables for "GATA2 Induces a Stem Cell-Like Transcriptional Program in Macrophages that Promotes Atherogenesis"

### Supplemental Materials

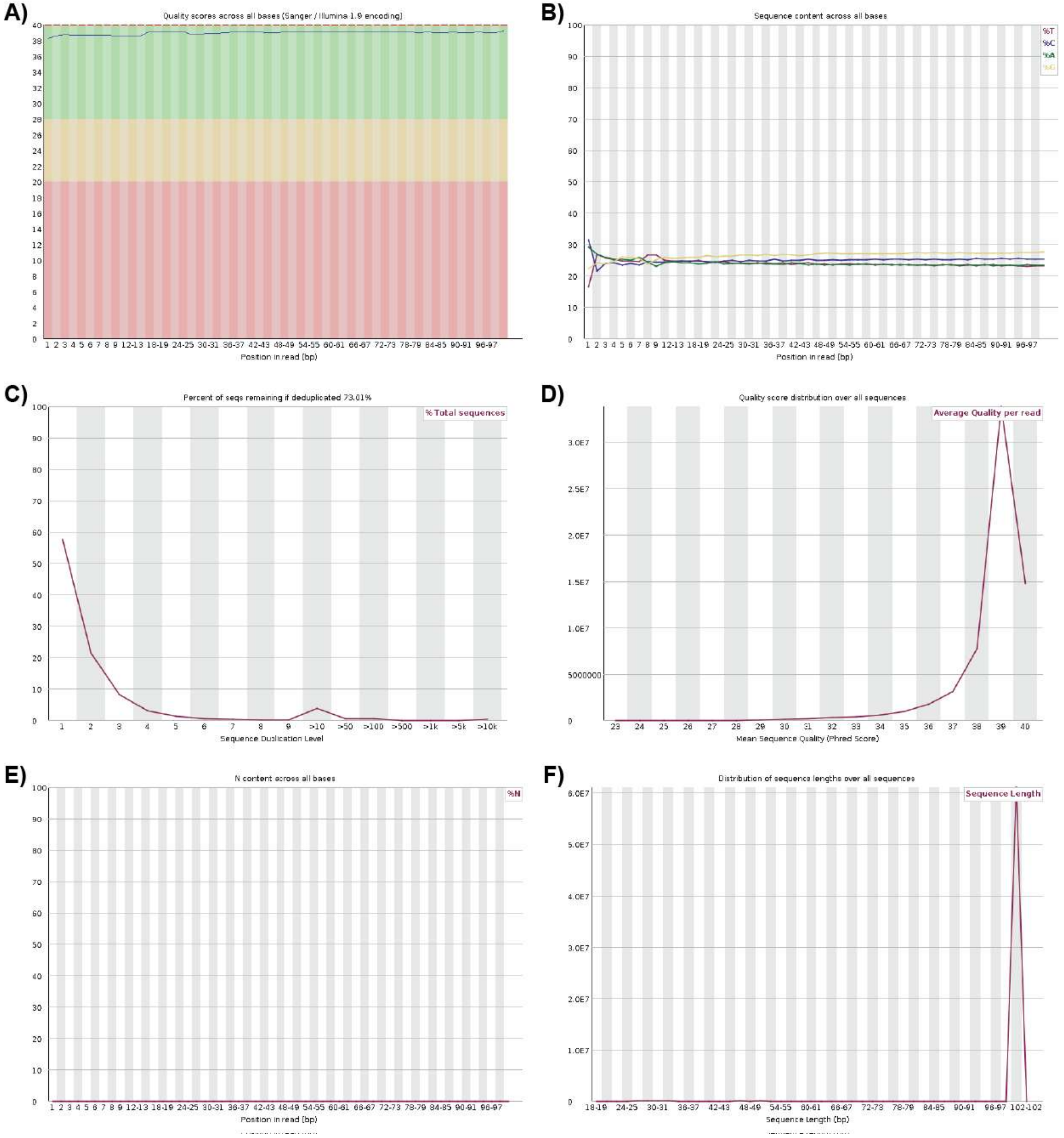


**Supplemental Figure 1: ChIP-seq quality controls. A)** Per-base mean quality scores (blue line) were >38 at all positions in the sequenced reads. **B)** The portion of the four DNA bases (A [green], T [red], C [blue], G [yellow]) are at approximately equal abundance at all positions in the sequenced reads. **C)** The percentage of read remaining after de-duplication was ~70%. **D)** Average quality scores across all reads. **E)** No 'N' contents were observed at any position in the sequence reads. **F)** Sequence length distribution of all reads after de-duplication. Data summarizes sequencing quality across two biological replicates.

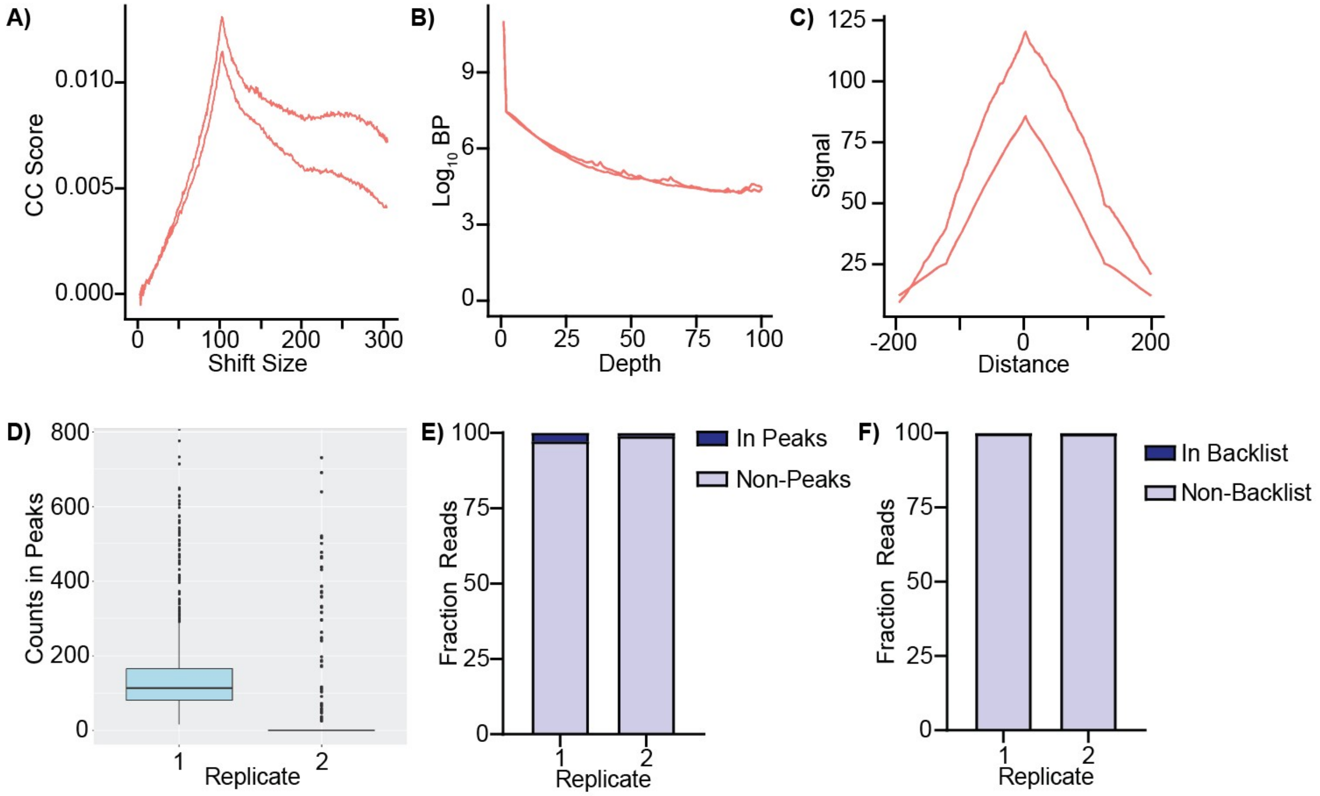


**Supplemental Figure 2: ChIP Distribution and Enrichment After Peak Calling. A-B)** ChIP signal distribution and structure as quantified by cross coverage (CC) score (A) and Log10 base pairs of genomes at differed read depths (B); each line quantifies a separate biological replicate. **C)** Confirmation of ChIP sequence enrichment within peaks, as quantified by the average signal profile; each line quantifies a separate biological replicate. **D)** The number of reads in peaks between the two biological replicates (Replicate 1/Replicate 2). **E)** Percentage of reads in enriched regions. **F)** Percentage of reads in blacklist regions.

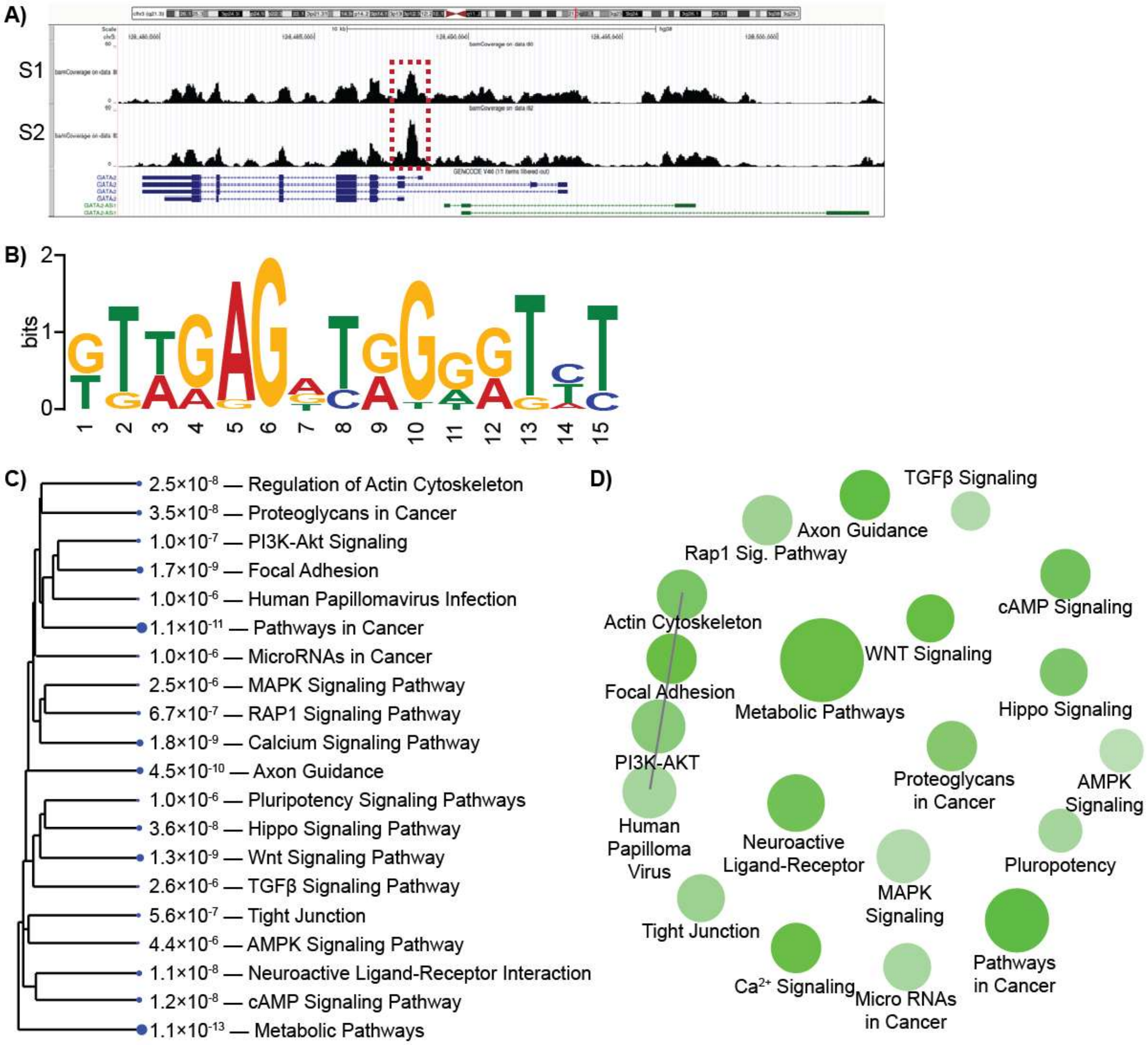


**Supplemental Figure 3: ChIP-seq Peak Calling and KEGG Pathways Correlations in Macrophages. A)** Alignment of GATA2 binding peaks from duplicate ChIP-seq samples (S1, S2) demonstrating an enrichment in GATA2 binding at the previously reported GATA2 binding site within the GATA2 gene (red box). **B)** MEME-ChIP analysis identification of the GATA2 binding motif. E-value = 4.7 × 10-7. **C)** A hierarchical clustering tree summarizes the correlation among KEGG pathways regulated by GATA2. Pathways with many shared genes are clustered together. Dots size correlates with statistical significance, numbers indicate p-value. **D)** KEGG analysis identified four nodes which were interconnected as they shared ≥20% of genes containing GATA2 binding peaks (line). All analyses are performed on replicate datasets.

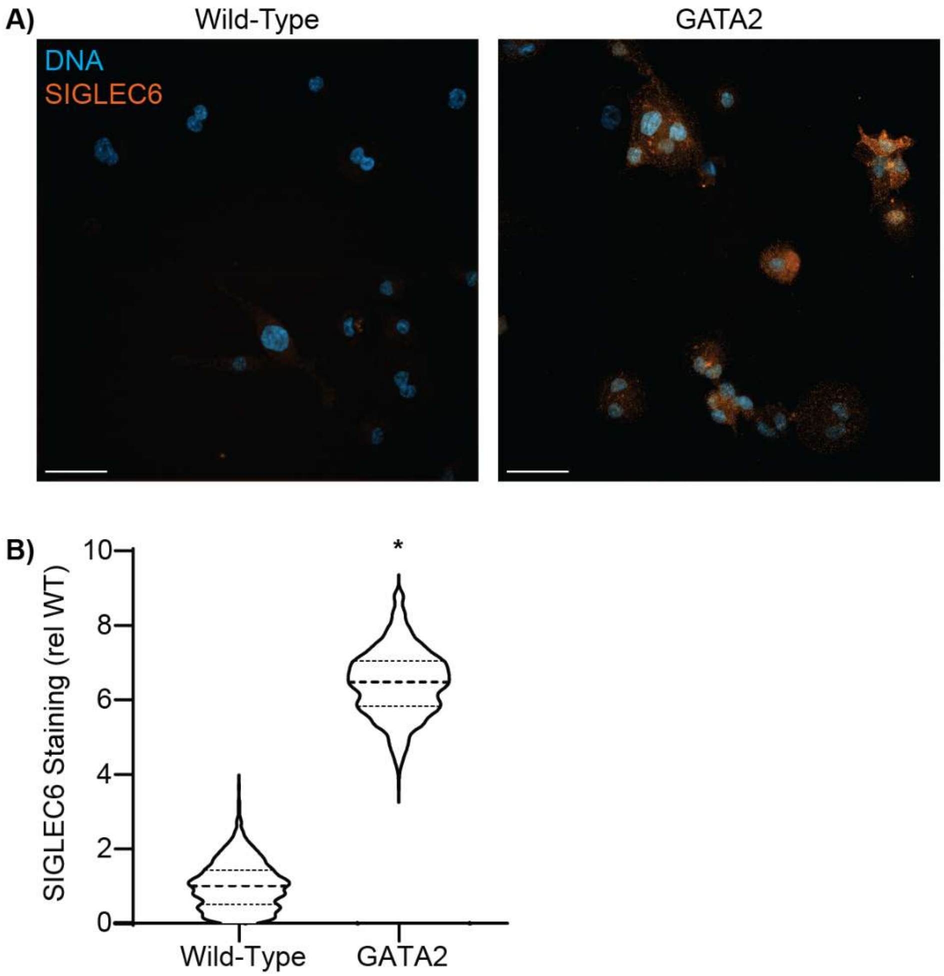


**Supplemental Figure 4: SIGLEC6 is a Marker of GATA-Expressing Macrophages. A)** Fluorescent micrographs of wild-type versus GATA2-overexpresing THP-1 macrophages stained for DNA (Hoechst, blue) and SIGLEC6 (orange). Both panels were imaged with identical staining and image acquisition parameters. Scale bars are 50 µm. **B)** Quantification of SIGLEC6 staining, normalized to the average intensity of SIGLEC6 staining on wild-type THP-1 macrophages. Data is plotted as the ensemble of at least 377 cells/group collected in 3 different experiments. * = $p < 0.05$ compared to wild-type, Students *T*-test.

**Supplemental Table 1: Promoter Regions Cloned for Dual-Luciferase Reporters.** All chromosome numbers and positions are based on the Human GRCh38/hg38 genome assembly. Start/End = starting and ending base pair positions of the cloned promoter, +/- indicate whether the gene is in the positive or negative strand of the chromosome.

| Promoter | Chromosome | Strand | Start | End | Size (bp) |
|---|---|---|---|---|---|
| GATA2 | 3 | - | 128,487,917 | 128,488,349 | 433 |
| ABCG1 | 21 | + | 42,209,416 | 42,210,730 | 1314 |
| DISP3 | 1 | + | 11,478,641 | 11,479,155 | 514 |
| IL-12B | 5 | - | 159,330,485 | 159,331,506 | 1021 |
| CCR2 | 3 | + | 46,353,585 | 46,354,117 | 566 |
| CIITA pI[1] | 16 | + | 10,876,893 | 10,877,241 | 348 |
| HLA-DRA[2] | 6 | + | 32,439,651 | 32,439,890 | 239 |

1. CIITA contains three promoters, of which, promoter 1 (pI) is active in myeloid cells[70].
2. MHC II allele which is representative of the regulation of MHC II.

**Supplemental Table 2: Antibodies Used in this Study**

| Antigen | Supplier | Clone | Purpose | Concentration |
|---|---|---|---|---|
| Human GATA2 | Abcam | EPR2822(2) | ChIP-seq | 50 µg/mL |
| Rabbit Isotype | ThermoFisher | 02-6102 | ChIP-seq isotype control | 50 µg/mL |
| Pan human-MHC II | Biolegend | Tu39 | Mouse-anti-Human opan-MHC II Immunostaining | 1:500 dilution |
| SIGLEC6 | R&D Systems | 767329 | SIGLEC6 immunostaining | 1:250 dilution |
| Anti-Mouse IgG Cy3 Fab | Jackson Immunoresearch | N/A | Secondary Fab fragment antibody | 10 µg/mL |
| Anti-Mouse IgG Alexa 647 Fab | Jackson Immunoresearch | N/A | Secondary Fab fragment antibody | 10 µg/mL |
| CellROX Orange | ThermoFisher Scientific | N/A | Reactive oxygen species staning | 5 µM |
| Hoechst 33342 | ThermoFisher Scientific | N/A | DNA staining | 5 µg/mL |

**Supplemental Table 3: Primer Sequences Used in this Study**

| Primer | Purpose | Sequence | Notes |
|---|---|---|---|
| GATA2-RACE | Gene-specific primer for 5' RACE | TTCGA ACCGC ATTAG AGTCG GGGTG CTGCG CATTC AGCAC G | Underlined region is homologous to GATA2 |
| GAPDH qPCR FWD | Forward qPCR primer | TCAAGGCTGAGAACGGGAAG | Tm = 57 |
| GAPDH qPCR REV | Reverse qPCR primer | CGCCCCACTTGATTTTGGAG | Tm = 57 |
| IL-12B qPCR FWD | Forward qPCR primer | GACATTCTGCGTTCAGGTCCAG | Tm = 60 |
| IL-12B qPCR REV | Reverse qPCR primer | CATTTTTGCGGCAGATGACCGTG | Tm = 61 |
